# SNF1-related protein kinase 2 directly regulate group C Raf-like protein kinases in abscisic acid signaling

**DOI:** 10.1101/2020.02.04.933978

**Authors:** Yoshiaki Kamiyama, Misaki Hirotani, Shinnosuke Ishikawa, Fuko Minegishi, Sotaro Katagiri, Fuminori Takahashi, Mika Nomoto, Kazuya Ishikawa, Yutaka Kodama, Yasuomi Tada, Daisuke Takezawa, Scott C. Peck, Kazuo Shinozaki, Taishi Umezawa

**Affiliations:** Graduate School of Bio-Applications and Systems Engineering, Tokyo University of Agriculture and Technology, Tokyo 184-8588, Japan.; Gene Discovery Research Group, RIKEN Center for Sustainable Resource Science, Ibaraki 305-0074, Japan.; Division of Biological Science, Nagoya University, Aichi 464-8602, Japan.; Center for Gene Research, Nagoya University, Aichi 464-8602, Japan.; Center for Bioscience Research and Education, Utsunomiya University, Tochigi 321-8505, Japan.; Graduate School of Science and Engineering, Saitama University, Saitama 338-8570, Japan.; Department of Biochemistry, University of Missouri, Columbia, MO 65211, USA.; Faculty of Agriculture, Tokyo University of Agriculture and Technology, Fuchu, Tokyo 183-8538, Japan.; PRESTO, Japan Science and Technology Agency, Saitama 332-0012, Japan.

## Abstract

A phytohormone abscisic acid (ABA) has a major role in abiotic stress responses in plants, and subclass III SNF1-related protein kinase 2 (SnRK2) mediates ABA signaling. In this study, we identified Raf36, a group C Raf-like protein kinase in Arabidopsis, as an interacting protein with SnRK2. A series of reverse genetic and biochemical analyses revealed that Raf36 negatively regulates ABA responses and is directly phosphorylated by SnRK2s. In addition, we found that Raf22, another C-type Raf-like kinase, functions partially redundantly with Raf36 to regulate ABA responses. Comparative phosphoproteomic analysis using Arabidopsis wild-type and *raf22raf36-1* plants identified proteins that are phosphorylated downstream of Raf36 and Raf22 *in planta*. Together, these results reveal a novel subsection of ABA-responsive phosphosignaling pathways branching from SnRK2.

## INTRODUCTION

Environmental stresses, such as drought, high salinity and low temperature, have adverse effects on plant growth and development. Abscisic acid (ABA) is a phytohormone that plays important roles in responses and adaptations to these stresses, as well as in embryo maturation and seed dormancy (Finkelstein, 2013; Shinozaki et al., 2003). The major ABA signaling pathway consists of three core components: ABA receptors, type 2C protein phosphatases (PP2Cs) and SNF1-related protein kinase 2s (SnRK2s) (Cutler et al., 2010; Umezawa et al., 2010). In this pathway, SnRK2s transmit ABA- or osmostress induced-signals thorough phosphorylation of downstream substrates, thereby promoting ABA- or stress-inducible gene expression and stomatal closure (Furihata et al., 2006; Geiger et al., 2009; Umezawa et al., 2013; Wang et al., 2013). The Arabidopsis genome contains 10 members of SnRK2, and they are classified into three subclasses (Hrabak et al., 2003; Yoshida et al., 2002). Among them, subclass III members, SRK2D/SnRK2.2, SRK2E/OST1/SnRK2.6 and SRK2I/SnRK2.3, are essential for ABA responses (Fujii and Zhu, 2009; Fujita et al., 2009; Nakashima et al., 2009; Umezawa et al., 2009).

Raf-like protein kinases were recently identified as regulators of ABA signaling. Among 80 putative MAPKKKs in Arabidopsis, 48 members are categorized as Raf-like subfamilies and can further be divided into 11 subgroups (B1-B4 and C1-C7) (Ichimura et al., 2002). In *Physcomitrella patens*, the *ARK* (also named *ANR* or *CTR1L*) gene is required for ABA-responsive SnRK2-activation, gene expression and drought, osmotic and freezing tolerance (Saruhashi et al., 2015; Stevenson et al., 2016; Yasumura et al., 2015). *ARK* encodes a B3 subgroup Raf-like protein kinase that phosphorylates SnRK2s *in vitro*, suggesting that ARK functions as a upstream kinase of SnRK2s (Saruhashi et al., 2015). In Arabidopsis, the B2 subgroup kinases Raf10 and Raf11 positively regulate seed dormancy and ABA responses by directly phosphorylating and activating subclass III SnRK2s (Lee et al., 2015; Nguyen et al., 2019). In addition to group B kinases, several group C Raf-like kinases have been associated with ABA responses. For example, Arabidopsis Raf43, a C5 kinase, promotes ABA sensitivity during seed germination and seedling root growth (Virk et al., 2015), whereas Raf22, a member of C6 subgroup, negatively regulates stress- or ABA-induced growth arrest (Hwang et al., 2018). However, despite these ABA-related genetic phenotypes, it is still unclear whether group C kinases directly regulate SnRK2-dependent signaling pathways.

In this study, we identified Raf36, a C5 group Raf-like kinase, as a protein that directly interacts with, and is phosphorylated by, SnRK2. Our evidence indicates that Raf36 functions as a negative regulator of ABA signaling pathway during post-germinative growth stage under the control of SnRK2. In addition, we revealed that Raf22, a C6 Raf-like kinase, functions partially redundantly with Raf36. Comparative phosphoproteomic analysis revealed that Raf36 and Raf22 are required for a subset of ABA-responsive phosphosignaling pathways. Collectively, unlike group B Rafs, which have been recently reported as an “accelerator” of ABA response upstream of SnRK2s, our results demonstrate that Arabidopsis group C Rafs, Raf22 and Raf36, functions as a “brake” of ABA response downstream of SnRK2s.

## RESULTS

### Raf36 interacts with subclass III SnRK2

To identify additional kinases that regulate ABA signaling pathways, we used the AlphaScreen^®^ assay to screen a collection of Arabidopsis MAPKKK proteins for their ability to physically interact with SRK2I (SnRK2.3), a subclass III SnRK2. From a pilot experiment, several Raf-like protein kinases were identified as candidate interactors with SRK2I (Supplemental Figure 1). Raf36, which belongs to a C5 subgroup kinase (Supplemental Figure 2), was one of the SRK2I-interacting proteins. Interaction between Raf36 and SRK2I, as well as between Raf36 and additional subclass III SnRK2s, SRK2D (SnRK2.2) and SRK2E (OST1/SnRK2.6), was confirmed by AlphaScreen^®^ assay (Figure 1A) and yeast two-hybrid assay (Figure 1B). SnRK2s were previously found within the cytosol and nuclei of Arabidopsis cells (Umezawa et al., 2009). We observed that Raf36-GFP is localized mainly in the cytosol (Figure 1C), and bimolecular fluorescence complementation (BiFC) assay confirmed that the interactions between SnRK2s and Raf36 take place in cytosol (Figure 1D) Together, these results demonstrate that Raf36 physically interacts with ABA-responsive SnRK2s both *in vitro* and *in vivo*.

**Figure 1.**
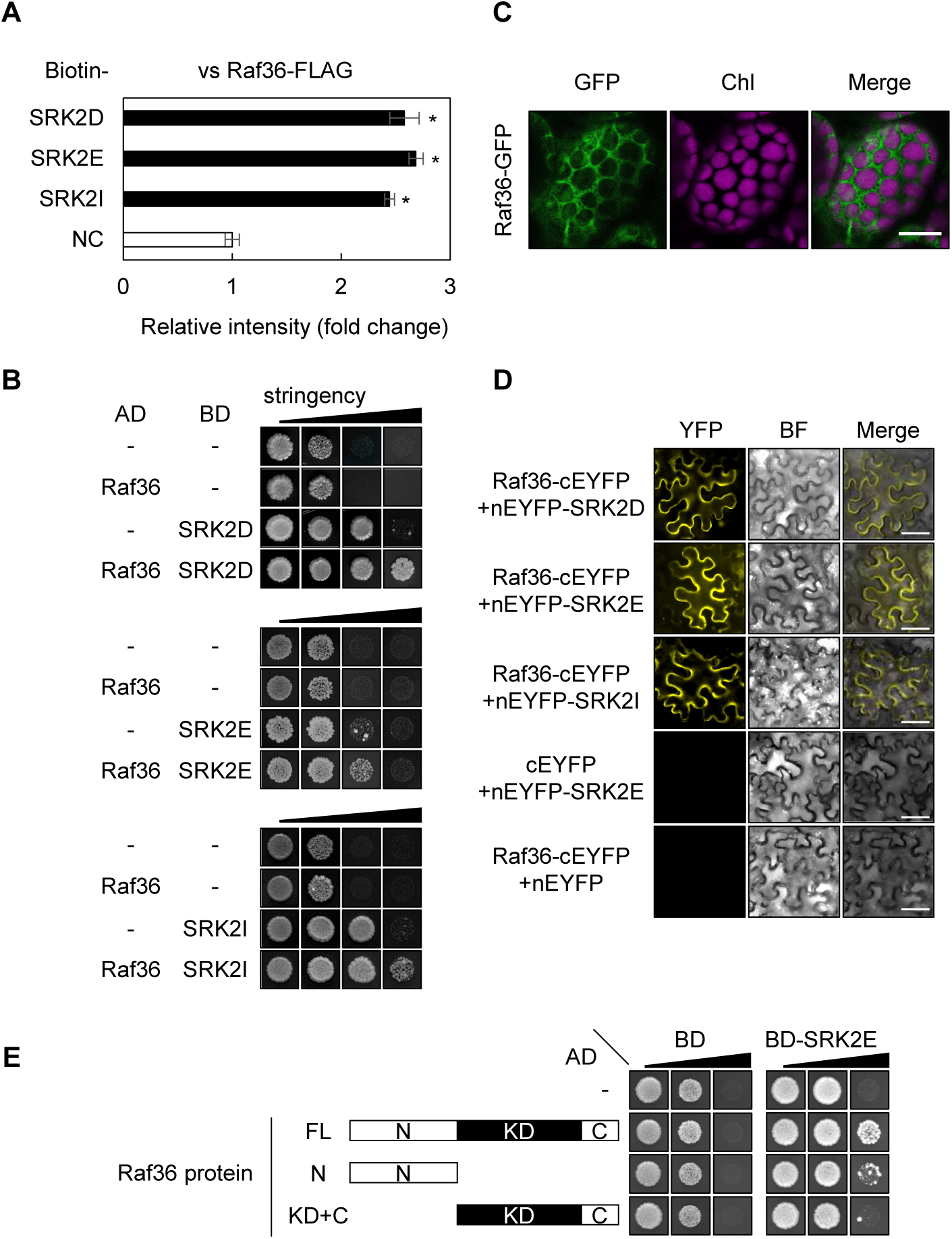
Raf36 interacts with subclass III SnRK2s. (A) AlphaScreen^®^ assay shows interaction of Raf36 and subclass III SnRK2s. Bars indicate means ± standard error (n=3), and asterisks indicate significant differences by Student’s *t* test (*P* < 0.05). (B) Yeast two-hybrid (Y2H) assay shows interaction between Raf36 and subclass III SnRK2s. Yeast cells expressing GAL4AD:Raf36 and GAL4BD:SnRK2s fusion proteins were incubated on SD media supplemented with or without 3-amino-1,2,4-triazole (3-AT) and lacking combinations of amino acids leucine (L), tryptophan (W) and histidine (H), as follows (in order from low to high stringency): −LW, −LWH, −LWH +10 mM 3-AT, −LWH +50 mM 3-AT. Photographs were taken at 10 days (SRK2D and SRK2E) or 12 days (SRK2I) after incubation. (C) Subcellular localization of Raf36-GFP in leaf mesophyll cells. Chl indicates chlorophyll autofluorescence. Scale bar, 20 µm. (D) BiFC assays for Raf36 and subclass III SnRK2s. SnRK2 and Raf36 were transiently expressed in *N. benthamiana* leaves by Agrobacterium infiltration. Empty vector constructs were used as negative controls. nEYFP and cEYFP represent the N- and C-terminal fragments of the EYFP, respectively. BF indicates bright field images. Scale bar, 50 µm. (E) Y2H assay for truncated versions of Raf36 and SRK2E. Yeast cells co-expressing GAL4AD:Raf36, Raf36 N or Raf36 KD+C and GAL4BD:SRK2E fusion proteins were incubated on SD media lacking L, W, H, and adenine (A), as follows (in order from low to high stringency): −LW, −LWH, −LWHA.

Next, we investigated which domain(s) of Raf36 may be responsible for the interaction with SnRK2s. According to the PROSITE database (https://prosite.expasy.org/), Raf36 contains an unknown N-terminal stretch (N, 1-206 aa), a predicted kinase catalytic domain (KD, 207-467 aa) and a short C-terminal domain (C, 468-525 aa) (Figure 1E). In yeast two-hybrid assay, SRK2E strongly interacted with Raf36 full-length protein (FL), but just slightly or not interacted with Raf36 N and KD+C alone, respectively (Figure 1E). These results indicated that the complete structure of Raf36 protein is required for the interaction with SnRK2.

### Raf36 negatively regulates ABA response at post-germination growth stage

To characterize the role of Raf36 in ABA signaling, we performed a series of functional analyses in Arabidopsis. Using qRT-PCR, we measured the abundance of *Raf36* mRNA in seedlings and detected a slight yet significant increase in *Raf36* transcripts after ABA treatment (Figure 2A). Next, we obtained two *Raf36* T-DNA insertion lines, GK-459C10 and SALK_044426C, designated as *raf36-1* and *raf36-2*, respectively (Supplemental Figure 3A). Using RT-PCR we confirmed the loss of *Raf36* transcripts in both mutants (Supplemental Figure 3B). We then measured rates of seed germination and cotyledon greening in the presence or absence of exogenous ABA. No difference in greening rate was observed between wild-type and mutant seedlings in the absence of ABA. However, with plants treated with 0.5 µM ABA, the greening rate of *raf36* mutants was significantly slower than wild-type (Figures 2B and 2C). This ABA-hypersensitive phenotype was complemented by *CaMV35S:Raf36-GFP* (Figure 2D and Supplemental Figure 3C). To assess if the delayed greening of *raf36* requires SnRK2 signaling, we generated a triple mutant, *raf36-1srk2dsrk2e*, and observed that it was less sensitive to ABA than *raf36-1* (Figures 2E and 2F). This result indicates that SRK2D and SRK2E are genetic modifiers of *raf36*-dependent ABA hyper-sensitivity in the greening response. Notably, seed germination rates were not significantly changed in either *raf36* mutant in the presence or absence of ABA (Supplemental Figure 3D). Taken together, our results suggested that Raf36 functions as a negative regulator of SnRK2-dependent ABA signaling during post-germinative growth.

**Figure 2.**
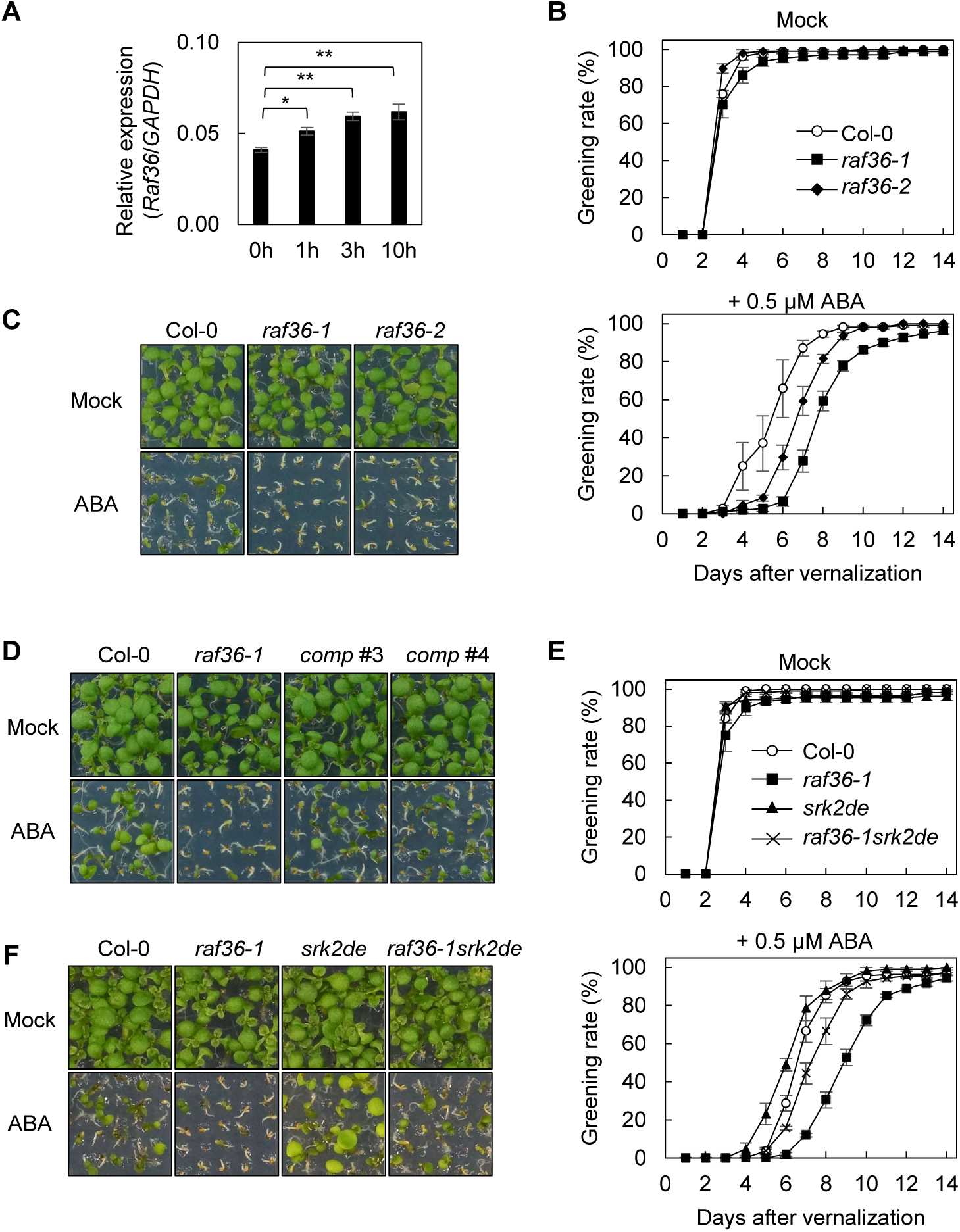
Raf36 negatively regulates SnRK2-dependent ABA response phenotypes in Arabidopsis seedlings. (A) Abundance of *Raf36* mRNA transcripts measured by quantitative RT-PCR. Total RNA was extracted from 1-week-old wild-type (WT) Col-0 Arabidopsis seedlings treated with 50 µM ABA for indicated periods. Bars indicate means ± standard error (n=3), and asterisks indicate significant differences by Student’s *t* test (\**P* < 0.05, \*\**P* < 0.01). (B and C) Quantification of the cotyledon greening rates of WT (Col-0), *raf36-1* and *raf36-2* on GM agar medium with or without 0.5 µM ABA. Data are means ± standard error (n=3). Each replicate contains 36 seeds. Photographs were taken 6 days after vernalization. (D) Functional complementation of *raf36-1* by *CaMV35S:Raf36-GFP*. Shown in photograph of seedlings grown for 7 days on GM agar medium in the presence or absence of 0.5 µM ABA. (E and F) Quantification of the cotyledon greening rates of WT (Col-0), *raf36-1*, *srk2dsrk2e* and *raf36-1srk2dsrk2e* on GM agar medium in the presence or absence of 0.5 µM ABA. Data are means ± standard error (n=3). Each replicate contains 36 seeds. Photographs were taken 6 days after vernalization.

### Raf36 is phosphorylated by SnRK2

To further examine the biochemical relationship between SnRK2 and Raf36, we prepared Raf36 and SRK2E recombinant proteins as maltose-binding protein (MBP)- or glutathione S-transferase (GST)-fusions. Raf36 protein auto-phosphorylated, demonstrating that Raf36 is an active kinase (Figure 3A). We found that Raf36 prefers Mn^2+^ for its kinase activity (Supplemental Figure 4A), as shown for other C-group Raf (Lamberti et al., 2011; Reddy and Rajasekharan, 2006; Rudrabhatla et al., 2006), and SRK2E prefers Mg^2+^ for its kinase activity (Supplemental Figure 4B). We then performed *in vitro* phosphorylation assays using kinase-dead forms of Raf36 and SRK2E as substrates. SRK2E phosphorylated Raf36 (K234N), while Raf36 did not phosphorylate SRK2E (K50N) (Figure 3A). This result suggested that Raf36 is a potential substrate of SnRK2, but not vice versa.

**Figure 3.**
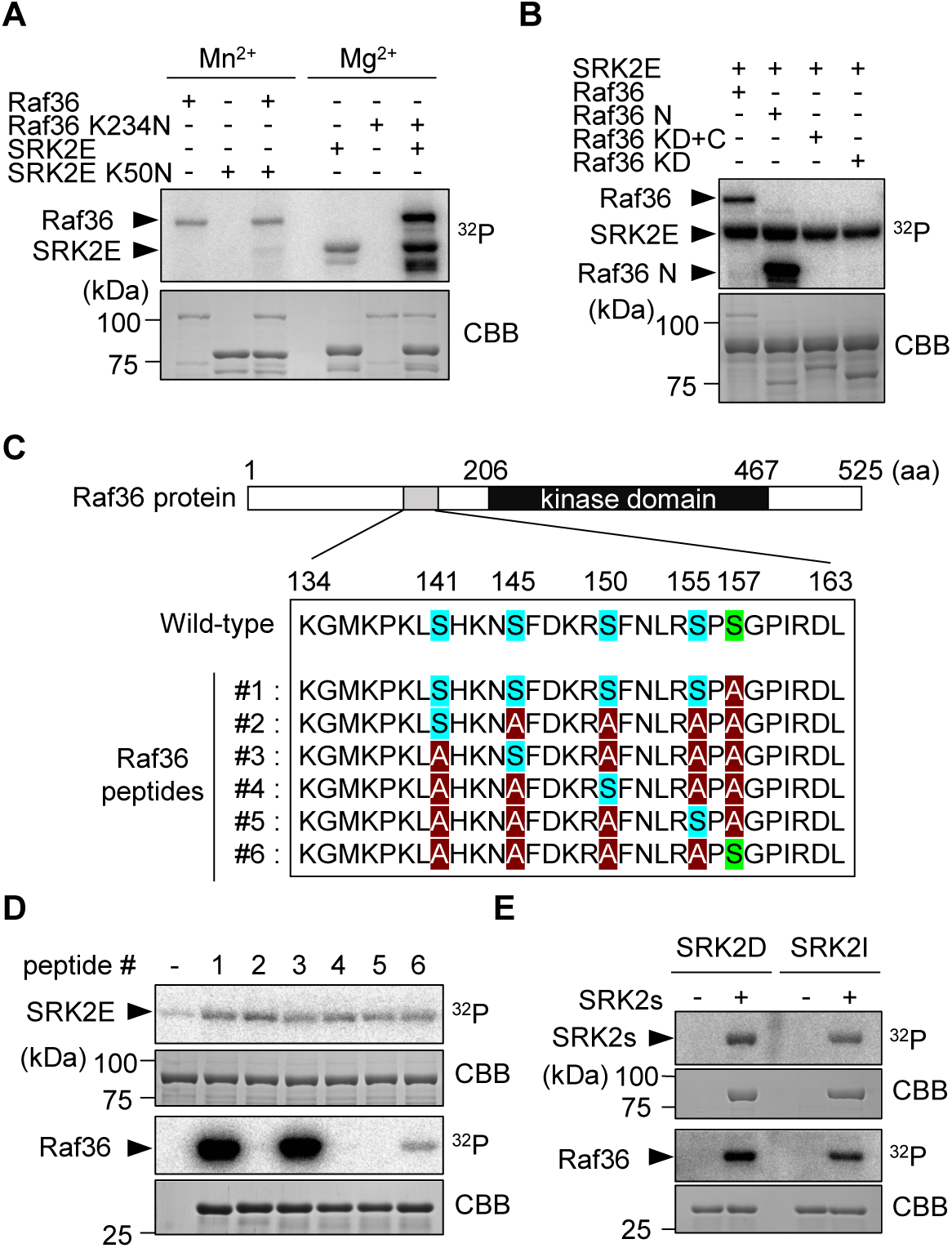
Subclass III SnRK2s directly phosphorylate Raf36. (A) *In vitro* phosphorylation assay using kinase-dead forms of GST-SRK2E (SRK2E K50N) or MBP-Raf36 (Raf36 K234N). Each kinase-dead form was co-incubated with an active GST-SRK2E or MBP-Raf36 kinase as indicated. Assays were performed in the presence of 5 mM Mn^2+^ (left 3 lanes) or 5 mM Mg^2+^ (right 3 lanes) with [γ-^32^P] ATP. (B) *In vitro* phosphorylation assay using truncated forms of MBP-tagged Raf36. Each MBP-Raf36 protein was incubated with MBP-SRK2E in the presence of 5 mM Mg^2+^ with [γ-^32^P] ATP. N: N-terminal region, KD: kinase domain, C: C-terminal region. (C) Schematic representation of six Raf36 (134-163) peptides tested as SRK2E substrates. Ser141, Ser145, Ser150 and Ser155 are labeled in blue, with alanine substitutions shown in red. Ser157, labeled in green, was replaced with alanine in Raf36 (134-163) peptides #1-#5. (D) *In vitro* phosphorylation of Raf36 peptides by MBP-SRK2E. (E) *In vitro* phosphorylation of GST-Raf36 (134-163) peptide #3 by MBP-SRK2D or MBP-SRK2I. Autoradiography (^32^P) and CBB staining (CBB) show protein phosphorylation and loading, respectively.

Additional *in vitro* kinase assays were performed using a series of truncated versions of Raf36 to identify the phosphorylation site(s) of Raf36. First, we observed that Raf36 proteins lacking the N-terminal region (Raf 36 KD+C and Raf36 KD) were not phosphorylated, suggesting this region is important for both auto-phosphorylation by Raf36 and trans-phosphorylation by SRK2E (Figure 3B). Second, Raf36 N (1-206) and Raf36 N (1-156) recombinant proteins were strongly phosphorylated, but Raf36 N (1-140) was slightly phosphorylated by SRK2E (Supplemental Figure 4C). These data suggested that the major phosphorylation site is located within 141-156 aa of the N-terminal region. Four serine (Ser) residues (i.e., Ser^141^, Ser^145^, Ser^150^ and Ser^155^) are present within this region. To identify which Ser residue(s) may be phosphorylated, we generated six peptides spanning this region of Raf36 (134-163). The peptides either contained all four serine residues (peptide #1), or had only a single Ser residue, with alanine substituted for the remaining Ser residues (peptides #2-#6) (Figure 3C). Ser^157^ was replaced with alanine in each peptide because it was outside of phosphorylated 141-156 aa region. Of these synthetic peptides, only peptides #1 and #3 were strongly phosphorylated by SRK2E, indicating Ser^145^ is the phosphorylation site (Figure 3D). SRK2D and SRK2I also phosphorylated Ser^145^ of Raf36 *in vitro* (Figure 3E). Taken together, these data show that subclass III SnRK2s phosphorylate Ser^145^ of Raf36.

### Raf22 functions partially redundantly with Raf36

We next tested if Raf kinases closely-related to Raf36 are also SnRK2 substrates. There are five kinases within Raf subgroups C5 and C6 (Figure 4A). Among them, HIGH LEAF TEMPERATURE 1 (HT1/ Raf19) functions independently of ABA (Hashimoto et al., 2006; Hashimoto-Sugimoto et al., 2016). Therefore, we focused our analyses on Raf43, Raf22 and Raf28. As shown in Figure 4B, SRK2E strongly phosphorylated Raf22, despite having no equivalent of Ser^145^ of Raf36 and sharing only 29 % identity with Raf36. In addition, SRK2E only weakly or not phosphorylated Raf43 and Raf28, respectively. We next tried to identify the phosphorylation site in Raf22. As described above, Raf28 was not phosphorylated by SnRK2, albeit with having 88 % identity with Raf22. Because SnRK2 kinases prefer [-(R/K)-x-x-(S/T)-] or [-(S/T)-x-x-x-x-(E/D)-] (Furihata et al., 2006; Umezawa et al., 2013), we searched the amino acid sequences of Raf22 and Raf28 for potential SnRK2 phosphorylation sites, and identified Ser^81^ within an [-R-H-Y-S-] motif in Raf22 that is converted to [-R-H-P-Y-S-] in Raf28. We introduced an alanine substitution at Ser^81^ in recombinant Raf22, and observed that this substitution nearly abolished phosphorylation by SRK2E (Figure 4C), indicating this serine residue is a SnRK2-phosphorylation site.

**Figure 4.**
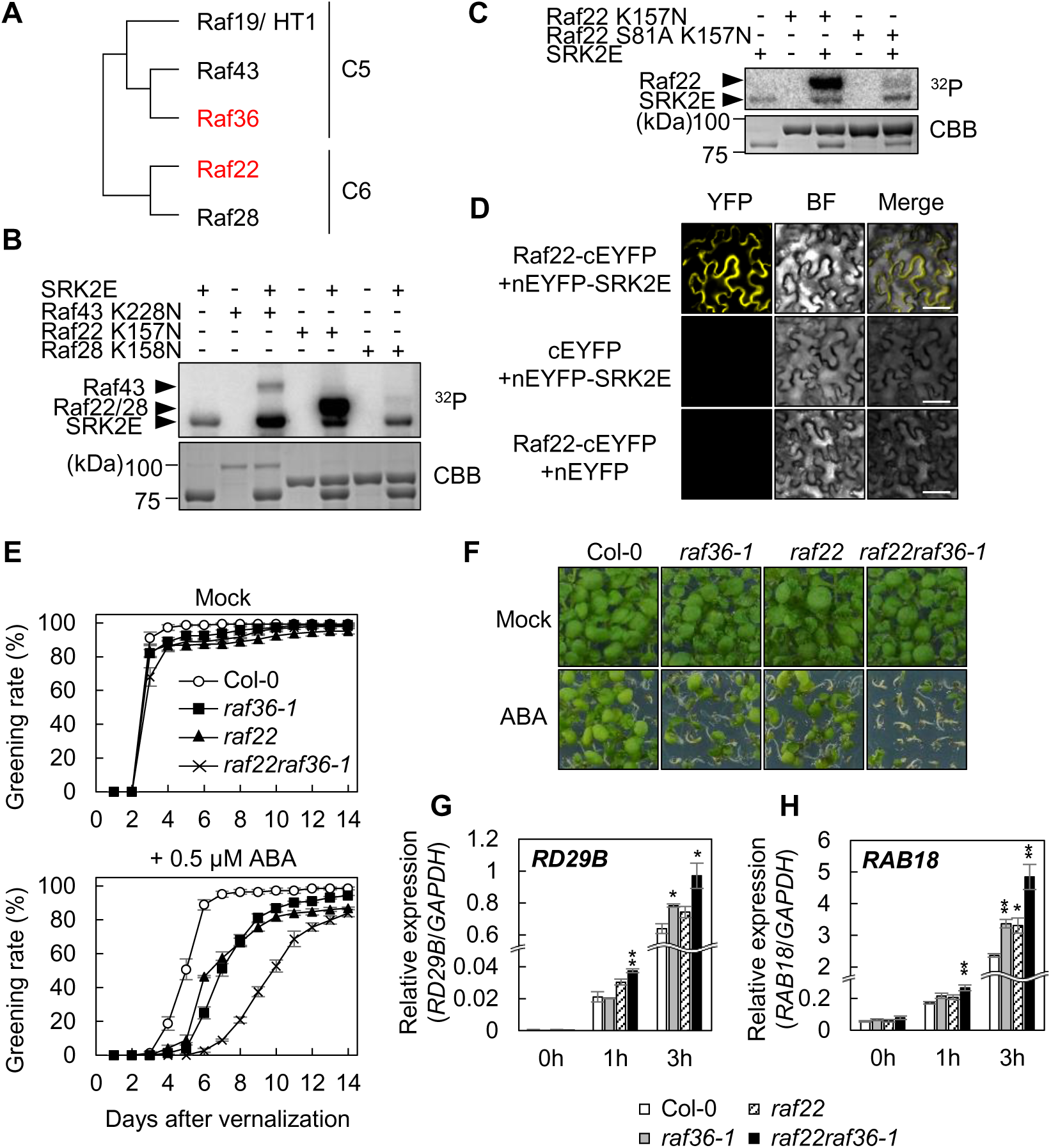
Raf22, a C6 Raf-like kinase, functions redundantly with Raf36. (A) Phylogenetic tree of subfamily C5 and C6 Raf-like kinases in Arabidopsis. (B) *In vitro* phosphorylation of C5/C6 Raf kinases by GST-tagged SRK2E. MBP-tagged kinase-dead forms of Raf43 (Raf43 K228N), Raf22 (Raf22 K157N) or Raf28 (Raf28 K158N) were used as substrates. (C) *In vitro* phosphorylation of kinase-dead Raf22 (K157N) and Raf22 (S81A K157N) proteins by GST-SRK2E. (D) BiFC assay of Raf22 and SRK2E in *N. benthamiana* leaves. nEYFP and cEYFP represent the N- and C-terminal fragments of the EYFP, respectively. BF indicates bright field images. Scale bar, 50 µm. (E and F) Quantification of the cotyledon greening rates of wild-type (Col-0), *raf36-1*, *raf22* and *raf22raf36-1* on GM agar medium with or without 0.5 µM ABA. Data are means ± standard error (n=4). Each replicate contains 36 seeds. Photographs were taken 9 days after vernalization. (G and H) Relative gene expression of ABA-responsive genes. Total RNA was extracted from 1-week-old plants including wild-type, *raf36-1*, *raf22* and *raf22raf36-1* treated with 50 µM ABA for indicated periods. Bars indicate means ± standard error (n=3) and asterisks indicate significant differences by Student’s *t* test (\**P* < 0.05, \*\**P* < 0.01).

Next, we functionally characterized the role of Raf22 in ABA-related phenotypes. Similar to *Raf36*, transcriptional level of *Raf22* was slightly up-regulated after exogenous ABA treatment (Supplemental Figure 5A). Using BiFC assay, we observed interaction between Raf22 and SnRK2s *in vivo* (Figure 4D and Supplemental Figure 5B). A T-DNA insertional mutant (SALK_105195C), *raf22*, showed a similar phenotype to *raf36*, i.e. ABA hypersensitivity in the post-germination growth (Figure 4E and 4F) but not in seed germination (Supplemental Figure 5C).

To test potential functional redundancy between Raf36 and Raf22, a *raf22raf36-1* double knockout mutant was generated. In the presence of ABA, *raf22raf36-1* showed a stronger ABA-hypersensitive phenotype relative to individual *raf22* and *raf36-1* mutants (Figures 4E and 4F). In addition, expression of ABA- and stress-responsive genes *RD29B* and *RAB18* were hyper-induced in *raf22raf36-1* seedlings (Figures 4G and 4H). Moreover, leaf water loss was examined because ABA also controls stomatal closure. However, leaf water loss of *raf22raf36-1* and individual *raf22* and *raf36-1* plants was similar to that of wild-type plants (Supplemental Figure 5D), suggesting that Raf36 and/or Raf22 has a minor role in stomatal movements. Taken together, these results demonstrated that Raf36 and Raf22 function redundantly in ABA signaling during post-germinative growth stage.

To examine whether the protein kinase activity of Raf36 and Raf22 are required for its negative regulation of ABA response, several complemented lines with kinase-dead form were generated. As shown in Figure 5A, the wild-type Raf36 complemented the ABA-hypersensitive phenotype of *raf36-1*, while the kinase-dead form of Raf36 (Raf36 K234N) could not. Similar results were also observed for *Raf22* (Supplemental Figure 6). These results suggested that the protein kinase activities of Raf36 and Raf22 are required for its function in ABA signaling.

**Figure 5.**
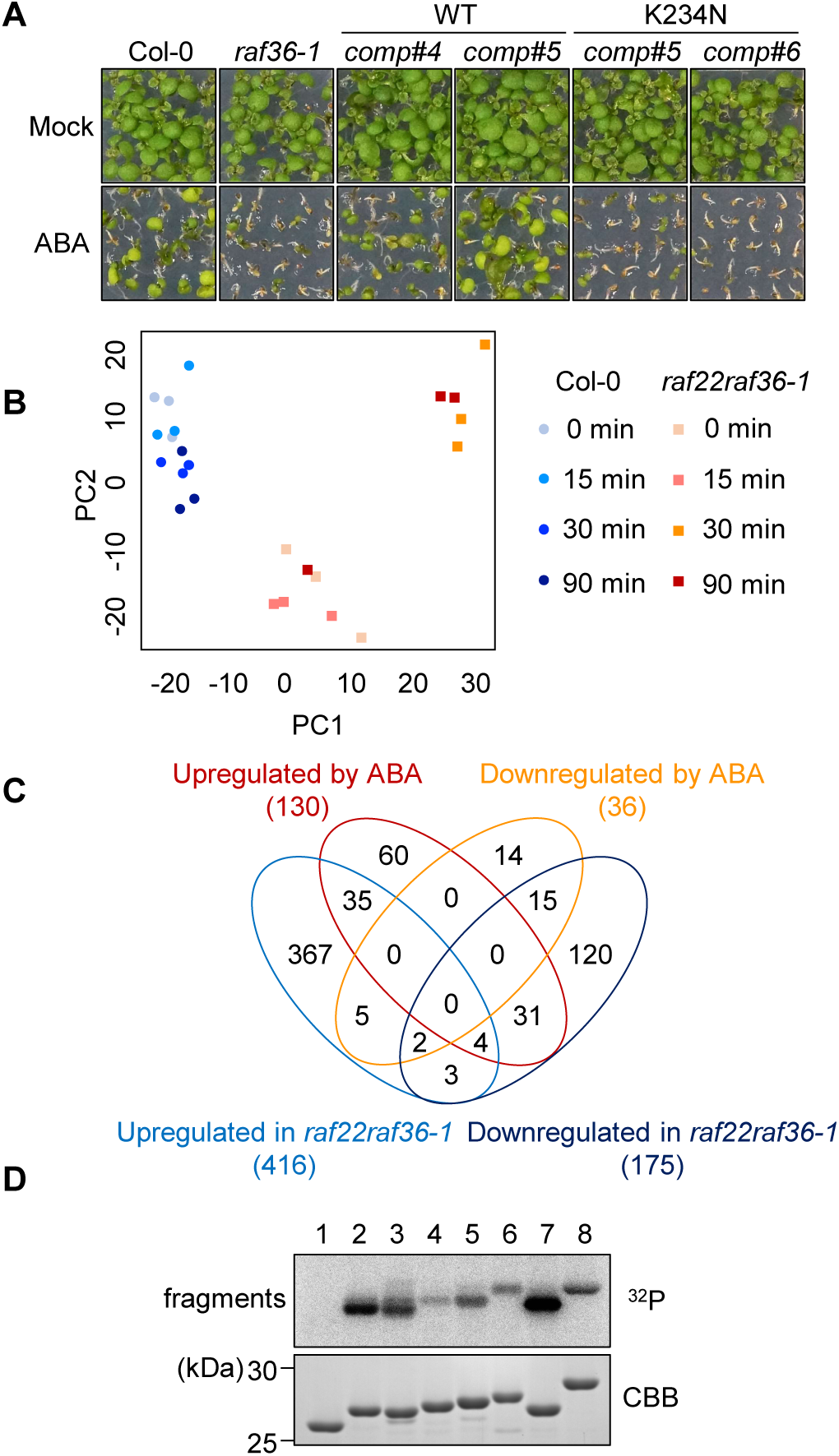
Phosphoproteomic analysis of wild-type and *raf22raf36-1* identifies ABA signaling components downstream of Raf kinases. (A) Functional complementation of *raf36-1* by *CaMV35S:Raf36-GFP* or *Raf36 K234N-GFP*. Shown in photograph of seedlings grown for 7 days on GM agar medium in the presence or absence of 0.5 µM ABA. (B) Principal component analysis of phosphoproteomic profiles of wild-type (WT) and *raf22raf36-1*. (C) Venn diagram of up- or down-regulated phosphopeptides in WT seedlings after 50 µM ABA treatment, and up- or down-regulated phosphopeptides in *raf22raf36-1* compared to WT (*P* < 0.05). (D) *In vitro* phosphorylation of GST-tagged phosphopeptides from proteins AT1G21630.1 (lane 3), AT1G20760.1 (lane 4), AT4G33050.3 (lane 5), AT1G60200.1 (lane 6), AT3G01540.2 (lane 7), AT1G20440.1 (lane 8) by MBP-Raf22. GST (lane 1) and GST-OLE1 fragment (143-173 aa, lane 2) were included as negative and positive controls, respectively. The autoradiography (^32^P) and CBB staining (CBB) show protein phosphorylation and loading, respectively.

### Raf36 and Raf22 regulate a subset of protein phosphorylation network in ABA response

To gain insight about ABA-responsive signaling pathway(s) that may be regulated by Raf36 and/or Raf22, we performed a comparative phosphoproteomic analysis of wild-type and *raf22raf36-1* seedlings treated with 50 µM ABA for 0, 15, 30 and 90 min. LC-MS/MS analysis identified a total of 1,500 phosphopeptides in both wild-type and the *raf22raf36-1* double mutant (Supplemental Dataset 1). Within this dataset 99% of identified phosphopeptides were singly phosphorylated (Supplemental Figure 7A). Phosphoserine, phosphothreonine, and phosphotyrosine accounted for 91.2%, 8.4% and 0.3% of phosphorylated residues, respectively (Supplemental Figure 7B). Consistent with the ABA-hypersensitive phenotype observed in the *raf22raf36-1* double mutant (Figures 4E and 4F), principal component analysis demonstrated that the phosphoproteome profiles of the ABA-treated *raf22raf36-1* mutant were different from that of ABA-treated wild-type (Figure 5B). In addition, the profiles of *raf22raf36-1* were significantly altered even before ABA treatment (Figure 5B).

Next, we identified phosphopeptides differentially regulated between wild-type and *raf22raf36-1*. ABA-responsive and Raf36 and/or Raf22 dependent phosphopeptides were screened by *P* value. After exogenous ABA treatment, 130 phosphopeptides were upregulated and 36 phosphopeptides were downregulated in wild-type plants (Figure 5C and Supplemental Dataset 2). In comparison with wild-type at each time point, a total of 416 and 175 phosphopeptides were upregulated and downregulated in r*af22raf36-1*, respectively (Figure 5C and Supplemental Dataset 2). Intriguingly, 53.8 % of ABA-upregulated phosphopeptides in wild-type were up- or down-regulated in *raf22raf36-1* double mutant, while 61.1 % of ABA-downregulated phosphopeptides in wild-type were up- or down-regulated in *raf22raf36-1*, indicating that large part of ABA-regulated phosphopeptides may be under the control of Raf36 and/or Raf22.

Subsets of phosphopeptides were further analyzed to evaluate potential differences in cellular responses to ABA between wild-type and *raf22raf36-1* plants. First, gene ontology (GO) analysis reported the term “response to abscisic acid” as significantly up- and down-regulated in the *raf22raf36-1* double mutant (Supplemental Figure 8C and 8D). In *raf22raf36-1*, RNA- and metabolism-related GO terms were significantly up- and down-regulated groups before ABA treatment, respectively (Supplemental Figures 9A and 9B). Second, we analyzed differentially-accumulating phosphopeptides for enrichment of motifs that correspond to kinase recognition sequences. Analysis using the algorithm motif-x (Wagih et al., 2016) identified two motifs, [-*p*(S/T)-P-] and [-(R/K)-x-x-*p*(S/T)-], that are MAPK- and SnRK2-/Calcium Dependent Protein Kinase (CDPK)-targeted sequences, respectively, as enriched in phosphopeptides from ABA-treated wild-type (Supplemental Figure 10). In addition, [-*p*(S/T)-P-] and [-(R/K)-x-x-*p*(S/T)-] containing phosphopeptides were also enriched in *raf22raf36-1* double mutant as compared to wild-type at each time point. Together, these two motifs account for over 70% of phosphopeptides that differentially accumulate in response to ABA in both wild-type and *raf22raf36-1*. This indicates Raf36 and Raf22 are directly or indirectly related to regulation of [-*p*(S/T)-P-] and/or [-(R/K)-x-x-*p*(S/T)-] *in vivo*.

To confirm our phosphoproteomic data, a subset of peptides were selected that were upregulated by ABA in wild-type but downregulated in *raf22raf36-1*. These candidates were synthesized as GST-fused 31 amino acid peptides and subjected to *in vitro* phosphorylation assay. The amino acids 143-173 of OLEOSIN1 (OLE1) was used as a positive control, because a previous study reported OLE1 as a substrate of Raf22 at seed stage (Ramachandiran et al., 2018). Consistent with our phosphorylation motif analysis (Supplemental Figure 10), both [-(R/K)-x-x-*p*(S/T)-] and [-*p*(S/T)-P-] containing phosphopeptides (AT1G21630.1: Calcium-binding EF hand family protein, AT1G20760.1: Calcium-binding EF hand family protein and AT4G33050.3: calmodulin-binding family protein EDA39 for [-(R/K)-x-x-*p*(S/T)-] motif; AT1G60200.1: splicing factor PWI domain-containing protein RBM25, AT3G01540.2: DEAD-box RNA helicase 1 (DRH1), AT1G20440.1: cold-regulated 47 (COR47) for [-*p*(S/T)-P-] motif) were directly phosphorylated by Raf22 (Figure 5D).

## DISCUSSION

SnRK2s are core components of abiotic stress signaling in plants, yet the full complement of SnRK2 substrates leading to responses necessary for stress tolerance remain unknown. To gain a better understanding of SnRK2-mediated signaling pathway(s), we aimed to identify signaling factors associated with SnRK2. In this study, Raf36 was identified as a candidate of SnRK2-interacting protein. Using several reverse genetic and biochemical analyses, we revealed that Raf36 and Raf22 are novel SnRK2 substrates which negatively regulate ABA responses in a partially redundant manner at post-germination growth stage through their protein kinase activities.

Our loss-of-function analyses revealed that Raf36 negatively regulates ABA responses (Figures 2B, 2C, 2D, 2E and 2F). Notably, loss of *Raf36* altered the rate of cotyledon greening, but did not affect seed germination or leaf water loss (Supplemental Figures 3D, 5C and 5D). These results suggest that Raf36 may function specifically in regulating ABA-induced post-germination growth arrest, a physiological response that may allow germinated seeds to survive under unfavorable conditions (Hwang et al., 2018). Raf36 is a member of subgroup C5 Raf-like MAPKKK family in Arabidopsis. Previous studies also reported that other group C kinases, Raf22 and Raf43, are associated with ABA responses, e.g. *raf22* mutant showed ABA-hypersensitive phenotype at post-germinative growth stage (Hwang et al., 2018) and *raf43* mutant showed ABA-hypersensitive phenotype at seed germination stage (Virk et al., 2015). Our results demonstrated that Raf36 and Raf22 functions partially redundantly in ABA-mediated post-germination growth arrest, because *raf22raf36-1* double knockout mutant showed a stronger ABA hypersensitivity than in the individual single mutants (Figures 4E and 4F). In addition to regulating post-germination growth arrest, our results show that Raf36 and Raf22 also regulate ABA-induced changes in gene expression and protein phosphorylation in older 1- or 2-week-old seedlings (Figures 4G, 4H, 5B, 5C, Supplemental Figures 8C and 8D). It has been known that some group C Raf-like kinases function in various tissues, e.g. HT1 or BLUE LIGHT-DEPENDENT H^+^-ATPASE PHOSPHORYLATION (BHP) in guard cells and Raf28 in embryogenesis (Hashimoto et al., 2006; Hayashi et al., 2017; Wang et al., 2018). Thus, Raf36 and Raf22 may also have different functions in different tissues and developmental stages.

In our experiments, kinase-dead forms of Raf36 or Raf22 did not complement phenotypes of *raf36-1* or *raf22* mutants, indicating that *in vivo* kinase activity of both proteins is necessary for regulating ABA responses (Figure 5A and Supplemental Figure 6). To identify possible substrates of Raf36 or Raf22, we performed a comparative phosphoproteomic analysis of ABA responses in wild-type and *raf22raf36-1* double mutant plants. Among phosphopeptides downregulated in *raf22raf36-1*, GO terms “Response to abscisic acid” or “Response to osmotic stress” were enriched (Supplemental Figure 8D), suggesting that Raf36 or Raf22 are involved in ABA-responsive phosphosignaling pathways. Consistent with this, *raf22raf36-1* double mutant showed enhanced salt tolerance compared to wild-type or the single mutant plants (Supplemental Figure 11), indicating that Raf36 and Raf22 are involved in ABA signaling at least under certain stress conditions. Functional analyses of detected phosphoproteins will be required for further understanding of phosphosignaling pathways under the control of Raf36 or Raf22. Unexpectedly, phosphoproteomic profiling revealed that Raf36 or Raf22 regulate some phosphosignaling pathways even in the absence of exogenous ABA treatment (Figure 5B). This suggests that Raf36 or Raf22 may have some roles under non-stressed conditions. A recent investigation proposed that basal ABA levels under well-watered conditions modulate plant metabolism and growth (Yoshida et al., 2019). Given that not only GO terms “Response to abscisic acid” but also metabolism-related GO terms were significantly enriched in ABA-nontreated *raf22raf36-1* mutant (Supplemental Figure 9B) and that *raf36* or *raf22raf36-1* showed a slight growth retardation in a normal condition (Supplemental Figure 12), Raf36 or Raf22 may regulate responses to basal ABA levels under normal conditions. Further analysis will be required to address this possibility in the future.

Several previous studies proposed the relationship between SnRK2 and Raf-like kinases, e.g. HT1 or PpARK (Hõrak et al., 2016; Matrosova et al., 2015; Saruhashi et al., 2015; Tian et al., 2015). Actually, PpARK, a subgroup B3 Raf, has been known as a direct upstream regulator of SnRK2, i.e. PpARK can activate SnRK2s by phosphorylating the activation loop of SnRK2 (Saruhashi et al., 2015). In this case, ARK acts as a positive regulator in ABA signaling in *Physcomitrella patens*. In addition, recent studies reported that Raf10, a subgroup B2 Raf, also acts as a positive regulator upstream of SnRK2s in Arabidopsis (Lee et al., 2015; Nguyen et al., 2019). In contrast to these results observed in group B Rafs, our results suggest that group C Rafs negatively regulate ABA signaling and that they are directly phosphorylated by SnRK2s. To clarify how they negatively regulate ABA responses, functional analysis of substrate candidates of Raf36 or Raf22, such as those from our phosphoproteomic data, will provide useful information for further understanding of group C Rafs in ABA signaling.

## METHODS

### Plant Materials

*Arabidopsis thaliana* ecotype Columbia (Col-0) was used as the wild-type. T-DNA insertion mutant lines, *raf36-1* (GK-459C10), *raf36-2* (SALK_044426C) and *raf22* (SALK_105195C) were obtained from ABRC or GABI-Kat. The *raf36-1raf22* double knockout mutant was generated by crossing *raf36-1* and *raf22*. The *SRK2E/OST1* knockout mutant, *srk2e* (SALK_008068), was used as described previously (Yoshida et al., 2002). The *srk2dsrk2e* double mutant was established by crossing *srk2d* (GABI-Kat 807G04) and *srk2e*, and the *raf36-1srk2dsrk2e* triple mutant was generated by crossing *srk2dsrk2e* double mutant and *raf36-1*. Seeds of wild-type, mutants or transgenic plants were sterilized, and sown on GM agar plates as described (Umezawa et al., 2009). After vernalization at 4 °C in the dark for 4 days, they were incubated in a growth chamber under a continuous light condition at 22 °C for indicated periods. To test ABA sensitivity, seeds were sown on GM agar medium with or without 0.5 µM ABA (Sigma, MO). Germination and greening rate were scored daily for 14 days according to previous study (Fujii et al., 2007).

### Plasmids

In this study, vector constructions were performed with Gateway cloning technology (Invitrogen, CA) unless otherwise noted. SRK2D, SRK2E and SRK2I cDNAs were previously cloned into pENTR/D-TOPO vector (Invitrogen, CA) (Umezawa et al., 2009). In addition, Raf36, Raf36 N (amino acid residues 1-206), Raf36 KD+C (207-525) and Raf22 cDNAs were cloned into pENTR/D-TOPO or pENTR1A vector and sequenced. Site-directed mutagenesis was carried out as previously described (Umezawa et al., 2013). Those cDNAs were transferred to destination vectors, such as pGBKT7 and pGADT7 (Takara Bio, Japan), pSITE-nEYFP-C1 and pSITE-cEYFP-N1 (ABRC), pGEX6p-2 (GE healthcare, IL) and pBE2113-GFP (Yoshida et al., 2002).

### Transgenic Plants

The pBE2113 vector, which drives C-terminal GFP-tagged Raf36 or Raf22 protein under the control of the Cauliflower Mosaic Virus (CaMV) 35S promoter, was constructed as described above. The transformation vector was introduced into *Agrobacterium tumefaciens* strain GV3101 by electroporation and transformed into Arabidopsis plants as described (Umezawa et al., 2004). The transformation lines were selected on GM agar medium containing 50 μg/ mL kanamycin and 200 μg/ mL claforan. Expression levels of transgene were checked by RT-PCR.

### AlphaScreen^®^

The AlphaScreen^®^ (Amplified Luminescent Proximity Homogeneous Assay) was carried out using an AlphaScreen^®^ FLAG^®^ (M2) Detection Kit (Perkin Elmer, MA) to detect protein-protein interactions. For screening of SnRK2-interacting MAPKKKs, 15 MAPKKKs were selected at random from the entire family members. Then, the C-terminal FLAG (DYKDDDDK)-tagged MAPKKK proteins were expressed in wheat germ extract (WGE) from *in vitro* synthesized mRNA obtained from PCR-amplified cDNAs (Nomoto and Tada, 2018). The N-terminal biotinylated SnRK2 proteins, such as SRK2D, E and I, were also synthesized in WGE. The protein quality (i.e., efficient synthesis with the expected molecular weight) of FLAG-tagged MAPKKKs and biotinylated SnRK2s was confirmed by western blotting with an anti-FLAG antibody and streptavidin, respectively. The FLAG-tagged MAPKKKs and biotinylated SnRK2s were mixed with acceptor beads coated with anti-FLAG antibody, donor beads coated with streptavidin, 0.01 % Tween-20 and 0.1 % bovine serum albumin (BSA) in sterilized water-diluted control buffer provided in the kit and then incubated at 21 °C for 12 h. The AlphaScreen^®^ luminescence was detected with the infinite^®^ M1000 Pro (TECAN, Switzerland). WGE with no expressed proteins was employed as negative control (NC) to estimate the luminescence caused by endogenous wheat germ proteins.

### Yeast two-hybrid analysis

Yeast two-hybrid analysis was employed using the MatchMaker GAL4 Two-Hybrid System 3 (Takara Bio, Japan) as previously described (Umezawa et al., 2009). *Saccharomyces cerevisiae* strain AH109 was co-transformed with various pairs of pGBKT7 vectors harboring SnRK2s (i.e., SRK2D, SRK2E and SRK2I) and pGADT7 vectors harboring Raf36. A single colony for each transformant grown on SD/-leucine (L)/-tryptophan (W) media was incubated in liquid media, and then evaluated on SD media supplemented with or without 3-amino-1,2,4-triazole (3-AT) and lacking combinations of amino acids leucine (L), tryptophan (W) and histidine (H), as follows: −LW, −LWH, −LWH +10 mM 3-AT, −LWH +50 mM 3-AT or on SD media lacking L, W, H, and adenine (A), as follows: −LW, −LWH, −LWHA. The plates were incubated at 30 °C for the optimal period.

### Microscopy analyses of fluorescent proteins

To perform *Agrobacterium*-mediated bimolecular fluorescence complementation (BiFC) assay, pSITE-nEYFP-C1 vectors harboring SnRK2s (i.e., SRK2D, SRK2E and SRK2I) or pSITE-cEYFP-N1 vectors harboring Raf-like kinases (i.e., Raf36 and Raf22) were introduced to *A. tumefaciens* strain GV3101(p19) by electroporation. A single colony for each transformant was cultured in LB media, and the media was substituted by 1/2 GM liquid media supplemented with 0.1 mM acetosyringone. SnRK2 and Raf transformants were mixed with various pairs, and then infiltrated into *Nicotiana benthamiana* leaves. Complemented YFP fluorescence of each samples was observed in epidermall cells of *N. benthamiana* at 3 days after infiltration with a fluorescence microscope BX53 (Olympus, Japan). For analysis of subcellular localization of Raf36-GFP, mesophyll cells of 2-week-old transgenic Arabidopsis plants expressing Raf36-GFP were observed with a confocal microscope SP8X (Leica Microsystems) with the time-gating method (Kodama, 2016), which completely eliminates chlorophyll autofluorescence when GFP imaging. GFP fluorescence was observed with 484 nm excitation and 494-545 nm emission with a gating time of 0.3–12.0 nsec. Chlorophyll autofluorescence was separately observed with 554 nm excitation and 640–729 nm emission.

### Preparation of Recombinant Proteins

DNA fragments of Raf36, Raf36 N (1-206), Raf36 KD+C (207-525), Raf36 KD (207-467), Raf36 N (1-156), Raf36 N (1-140), Raf22, Raf43, Raf28, SRK2D, SRK2E and SRK2I were amplified from cDNAs, and they were fused in-frame to pMAL-c5X vector (New England Biolabs, MA). Amino acid substitutions, such as Raf36 K234N, Raf22 K157N, Raf22 S81A K157N, Raf43 K228N, Raf28 K158N and SRK2E K50N, were introduced by site-directed mutagenesis. The MBP-fusion proteins were expressed and purified from *E. coli* BL21 (DE3) using Amylose Resin (New England Biolabs) according to the manufacturer’s instructions. GST-SRK2E and GST-SRK2E K50N proteins were expressed using pGEX6p-2 (GE healthcare, IL) and purified using Glutathione Sepharose 4B resin (GE healthcare, IL). Both MBP-tagged and GST-tagged recombinant proteins were further purified with a Nanosep^®^ 30-kDa size-exclusion column (PALL, NY). In addition, to identify phosphorylation sites in Raf36, six types of 30-amino-acid peptides with mutations were designed as Raf36 peptides (134-163) #1-#6 as shown in Fig. 3C. Similarly, to confirm Raf22-dependent phosphorylation, six phosphopeptides and OLE1 as a positive control, were designed as 31-amino-acid peptides including the putative phosphorylation site(s). These peptides were expressed and purified from *E. coli* BL21 (DE3) using pGEX4T-3 vector (GE healthcare, IL).

### *In vitro* phosphorylation assay

*In vitro* phosphorylation assays were performed as described previously with some modifications (Umezawa et al., 2009). The recombinant proteins of SnRK2, Raf or substrates were mixed with indicated pair(s) and incubated in 50 mM Tris-HCl (pH 7.5), 5 mM MgCl_2_ or 5 mM MnCl_2_, 50 µM ATP and 0.037 MBq of [γ-^32^P] ATP (PerkinElmer, MA) at 30 °C for 30 min. Samples were subsequently separated by SDS-PAGE, and phosphorylation levels were detected by autoradiography with BAS-5000 (Fujifilm, Japan).

### Water loss analysis

To measure leaf water loss, 2-week-old seedlings were transferred from GM agar medium to soil, and the plants were grown under a 16 h/8 h (light/dark) photoperiod at 22 °C for another 2 weeks. The fully expanded rosette leaves were detached from 4- to 5-week-old plants and placed on weighing dishes. These dishes were kept under the same conditions used for seedling growth on soil, and then their fresh weights were monitored at the indicated times with three replicates per time-point. One replicate consists of 5 individual leaves. Water loss was calculated as a percentage of relative weight at the indicated times versus initial fresh weight.

### RNA extraction and qRT-PCR

For quantitative reverse transcription PCR (qRT–PCR) analysis, total RNA was extracted by LiCl precipitation from 1-week-old seedlings treated with 50 µM ABA for indicated periods. 1 µg of total RNA treated with RNase-free DNase I (Nippon Gene, Japan) was used for reverse transcription with ReverTra Ace^®^ reverse transcriptase (TOYOBO, Japan). qRT-PCR analysis was performed using GoTaq^®^ qPCR Master Mix (Promega, WI) with Light Cycler 96 (Roche Life Science, CA). For normalization, *GAPDH* was used as an internal control. The gene-specific primers used for qRT-PCR analysis were shown in Dataset S5.

### Phosphoproteomic analysis

Following imbibition with 50 µM ABA, 2-week-old Arabidopsis seedling of wild-type (Col-0) and *raf22raf36-1* were used for phosphoproteomic analysis with three biological replicates. Total crude protein was extracted from grounded sample. The phosphoproteomic analyses were performed as previously described with 400 µg of total crude protein (Ishikawa et al., 2019a, 2019b; Nakagami et al., 2010; Sugiyama et al., 2007; Umezawa et al., 2013). Enriched phosphopeptides by using HAMMOC method (Sugiyama et al., 2007) were analyzed with a LC-MS/MS system, TripleTOF 5600 (AB-SCIEX). Peptides and proteins were identified using the database (TAIR 10) with Mascot (Matrix Science, version 2.4.0). The false discovery score was calculated using Benjamini-Hochberg method and set to 5 %. Each phosphorylation site was assessed by the site localization probability score calculated with the mascot delta score and defined confident as > 0.75 (Ishikawa et al., 2019a). Skyline version 4.2 (Maccoss lab software) was used for quantification of phosphopeptides on the basis of LC-MS peak area. All raw data files were deposited in the Japan Proteome Standard Repository Database (jPOST; JPST000630, https://repository.jpostdb.org/preview/20386920535e1580015868a, access key; 3457). Each phosphoproteomic sample was compared by principal component analysis by using all identified phosphopeptides and their phosphorylation level. The motif analysis was conducted using the Motif-X algorithm. DAVID (https://david.ncifcrf.gov) and REViGO (http://revigo.irb.hr) were used for GO analysis.

### Accession numbers

Sequence data from this article can be found in the GenBank/EMBL data libraries under the following accession numbers: *SRK2D, AT3G50500; SRK2E, AT4G33950, SRK2I, AT5G66880, Raf36, AT5G58950; Raf22, AT2G24360; Raf43, AT3G46930; Raf28, AT4G31170; RD29B, AT5G52300*, *RAB18, AT5G66400* and *OLE1, AT4G25140*.

## Supplemental Information

**Supplemental Figure 1.** AlphaScreen^®^ assay for screening of SnRK2-interacting MAPKKKs.

**Supplemental Figure 2.** A phylogenetic tree of Arabidopsis Raf-like kinases.

**Supplemental Figure 3.** Isolation and characterization of *raf36-1* and *raf36-2* T-DNA insertion lines.

**Supplemental Figure 4.** *in vitro* phosphorylation assays determining preference for Mg^2+^ or Mn^2+^.

**Supplemental Figure 5.** The characterization of Raf22 in ABA response.

**Supplemental Figure 6.** Raf22 protein kinase activity is required for its function in ABA signaling.

**Supplemental Figure 7.** An overview of phosphoproteomic analysis.

**Supplemental Figure 8.** GO analysis of phosphopeptides in wild-type and *raf22raf36-1*.

**Supplemental Figure 9.** GO analysis of phosphopeptides in *raf22raf36-1* in normal condition.

**Supplemental Figure 10.** Motif analysis of phosphopeptides in wild-type and *raf22raf36-1*.

**Supplemental Figure 11.** Salt tolerance of *raf22raf36-1* plants.

**Supplemental Figure 12.** Dwarf phenotype of *raf36* and *raf22raf36-1* plants under normal condition.

**Supplemental Dataset 1.** List of phosphopeptides detected in this study.

**Supplemental Dataset 2.** List of responded phosphopeptides during ABA treatment.

**Supplemental Dataset 3.** List of GO terms for phosphopeptides in this study.

**Supplemental Dataset 4.** Classification of phosphopeptides by motif groups.

**Supplemental Dataset 5.** Primer sequences used for quantitative RT-PCR (qRT-PCR).

## Acknowledgments

We thank Dr. Yoichi Sakata (Tokyo University of Agriculture, Japan), Dr. José M. Barrero (CSIRO, Australia) and Dr. Paul E. Verslues (Academia Sinica, Taiwan) for valuable discussion. We also thank Tomotaka Itaya, Ryo Yoshimura (Nagoya University, Japan) and Mrs. Saho Mizukado (RIKEN, Japan) for their expert technical assistance. We are grateful to ABRC and GABI-Kat project for providing Arabidopsis T-DNA insertional mutants. This work was partly supported by the Japan Society for the Promotion of Science (JSPS) KAKENHI [JP15H04383, 16KK0160, 19H03240] to T.U., and JST PRESTO [P13413773] to T.U.

## Author contributions

Yo.K. and T.U. designed research; Yo.K., M.H., S.I., F.M. and S.K. performed research; F.T., M.N., K.I., Yu.K., Y.T., D.T. and K.S. contributed new reagents/analysis tools; Yo.K., S.I., F.M. and S.K. analyzed data; and Yo.K., S.C.P. and T.U. wrote the paper.

## Figure Legends

**Supplemental Figure 1. AlphaScreen^®^ assay for screening of SnRK2-interacting MAPKKKs.**

Interaction of SnRK2 with MAPKKKs was tested by AlphaScreen^®^ assay. N-terminal biotin-tagged SRK2I protein and C-terminal FLAG-tagged MAPKKK proteins were synthesized by *in vitro* translation system in wheat germ extracts. After incubating in a reaction buffer including AlphaScreen^®^ acceptor and donor beads for 12 h, the AlphaScreen^®^ luminescence intensity was analyzed with a multi-mode plate reader (TECAN M1000pro).

**Supplemental Figure 2. A phylogenetic tree of Arabidopsis Raf-like kinases.**

Amino acid sequences of predicted kinase domains from Raf-like kinases were aligned using ClustalW. The phylogenetic tree was generated using MEGA-X software with the neighbor-joining method. The Raf-like kinases were classified as B1-B4 and C1-C7 subfamilies, according to Ichimura et al., 2002. The C5 subfamily is shown in bold red letters.

**Supplemental Figure 3. Isolation and characterization of *raf36-1* and *raf36-2* T-DNA insertion lines.**

(A) Schematic depiction of *Raf36* genomic DNA with T-DNA insertions. Black boxes and lines indicate exons and introns, respectively. (B) RT-PCR analysis of *Raf36* transcript levels in wild-type (Col-0), *raf36-1* and *raf36-2* seedlings. *GAPDH* was used as a positive control. (C) RT-PCR analysis of *Raf36* transcript levels in *raf36-1* and complementation lines (*comp*#3 and *comp*#4). *GAPDH* was used as a positive control. (D) Germination rates of wild-type (Col-0), *raf36-1* and *raf36-2* on GM agar medium in the presence or absence of 0.5 µM ABA. Data are means ± standard error (n=3). Each replicate contains 36 seeds.

**Supplemental Figure 4. *in vitro* phosphorylation assays determining preference for Mg^2+^ or Mn^2+^.**

(A) Effects of Mg^2+^ or Mn^2+^ on kinase activity of Raf36. MBP-tagged Raf36 and α-casein were incubated with [γ-^32^P] ATP in the presence of 5 mM Mg^2+^ (left) or 5 mM Mn^2+^ (right). Autophosphorylation (upper) and substrate phosphorylation (lower) signals were visualized by autoradiography. Coomassie Brilliant Blue (CBB) staining shows protein loading. (B) Effects of Mg^2+^ or Mn^2+^ on kinase activity of SRK2E. MBP-tagged SRK2E and histone as substrate were co-incubated with [γ-^32^P] ATP in the presence of 5 mM Mg^2+^ (left) or 5 mM Mn^2+^ (right). Autophosphorylation (upper) and substrate phosphorylation (lower) signals were visualized by autoradiography. Coomassie Brilliant Blue (CBB) staining shows protein loading. (C) *In vitro* phosphorylation assay to identify SnRK2-phosphorylation site(s) in the N-terminal region of Raf36.

**Supplemental Figure 5. The characterization of Raf22 in ABA response.**

(A) qRT-PCR analysis of *Raf22* mRNA transcript abundance. Total RNA was extracted from 1-week-old wild-type seedlings treated with 50 µM ABA for indicated periods. Bars indicate means ± standard error (n=3). Asterisks showed significant differences by Student’s *t* test (\**P* < 0.05, \*\**P* < 0.01). (B) BiFC assays of Raf22 and SnRK2s. SRK2D or SRK2I were transiently expressed with Raf22 in *N. benthamiana* leaves by Agrobacterium infiltration. nEYFP and cEYFP represent the N- and C-terminal fragments of the EYFP, respectively. BF indicates bright field images. Scale bar, 50 µm. (C) Germination rates of wild-type (Col-0), *raf36-1*, *raf22* and *raf22raf36-1* on GM agar medium with/without 0.5 µM ABA. Data are means ± standard error (n=3). Each replicate contains 36 seeds. (D) Water loss from detached leaves of wild-type (Col-0), *raf36-1*, *raf22* and *raf22raf36-1* plants. The *srk2e* mutant was included as a positive control. Data are means ± standard error (n=3). Each replicate consists of five individual leaves.

**Supplemental Figure 6. Raf22 protein kinase activity is required for its function in ABA signaling.**

Functional complementation of *raf22* mutant by introducing *Raf22-GFP* or *Raf22 K157N-GFP*. Photographs were taken 7 days after vernalization.

**Supplemental Figure 7. An overview of phosphoproteomic analysis.**

(A) Distribution of the number of phosphosites per peptide. (B) Distribution of phosphorylated residues in each peptide. *p*S, *p*T and *p*Y indicate phospho-serine, phospho-threonine and phospho-tyrosine, respectively.

**Supplemental Figure 8. GO analysis of phosphopeptides in wild-type and *raf22raf36-1*.**

Each graph represents GO terms for upregulated (A) and downregulated phosphopeptides (B) in ABA-treatment wild-type (WT) seedlings, or GO terms of up- (C) or down-regulated phosphopeptides (D) in *raf22raf36-1* as compared with WT. GO terms were evaluated by DAVID program and visualized with REVIGO (*P* < 0.05). Circle color and size show *P* value and frequency (%), respectively.

**Supplemental Figure 9. GO analysis of phosphopeptides in *raf22raf36-1* in normal condition.**

Each graph represents GO terms for phosphopeptides up- (A) or down-regulated (B) in *raf22raf36-1* under normal condition (*P* < 0.05). GO terms were evaluated by DAVID program and visualized with REVIGO.

**Supplemental Figure 10. Motif analysis of phosphopeptides in wild-type and *raf22raf36-1*.**

Phosphorylation motifs in up- or down-regulated phosphopeptides in response to ABA in wild-type seedlings, and motifs in up- or down-regulated phosphopeptides in *raf22raf36-1* in comparison with wild-type.

**Supplemental Figure 11. Salt tolerance of *raf22raf36-1* plants.**

Wild-type (Col-0), *raf36-1*, *raf22* and *raf22raf36-1* seeds were germinated on GM agar medium with or without 150 mM NaCl. Photographs were taken 19 days after vernalization.

**Supplemental Figure 12. Dwarf phenotype of *raf36* and *raf22raf36-1* plants under normal condition.**

Wild-type, *raf22*, *raf36* and *raf22raf36-1* plants grown at 22°C under 16/8 h photoperiod for 29 days. Scale bar, 3 cm.

## Parsed Citations

Cutler, S.R., Rodriguez, P.L., Finkelstein, R.R., and Abrams, S.R. (2010). Abscisic Acid: Emergence of a Core Signaling Network. Annu. Rev. Plant Biol. 61: 651–679.

Finkelstein, R. (2013). Abscisic Acid Synthesis and Response. Arab. Book Am. Soc. Plant Biol. 11.

Fujii, H., Verslues, P.E., and Zhu, J.-K. (2007). Identification of Two Protein Kinases Required for Abscisic Acid Regulation of Seed Germination, Root Growth, and Gene Expression in Arabidopsis. Plant Cell 19: 485–494.

Fujii, H. and Zhu, J.-K. (2009). Arabidopsis mutant deficient in 3 abscisic acid-activated protein kinases reveals critical roles in growth, reproduction, and stress. Proc. Natl. Acad. Sci. 106: 8380–8385.

Fujita, Y. et al. (2009). Three SnRK2 Protein Kinases are the Main Positive Regulators of Abscisic Acid Signaling in Response to Water Stress in Arabidopsis. Plant Cell Physiol. 50: 2123–2132.

Furihata, T., Maruyama, K., Fujita, Y., Umezawa, T., Yoshida, R., Shinozaki, K., and Yamaguchi-Shinozaki, K. (2006). Abscisic acid-dependent multisite phosphorylation regulates the activity of a transcription activator AREB1. Proc. Natl. Acad. Sci. 103: 1988–1993.

Geiger, D., Scherzer, S., Mumm, P., Stange, A., Marten, I., Bauer, H., Ache, P., Matschi, S., Liese, A., Al-Rasheid, K.A.S., Romeis, T., and Hedrich, R. (2009). Activity of guard cell anion channel SLAC1 is controlled by drought-stress signaling kinase-phosphatase pair. Proc. Natl. Acad. Sci. 106: 21425–21430.

Hashimoto, M., Negi, J., Young, J., Israelsson, M., Schroeder, J.I., and Iba, K. (2006). Arabidopsis HT1 kinase controls stomatal movements in response to CO2. Nat. Cell Biol. 8: 391–397.

Hashimoto-Sugimoto, M., Negi, J., Monda, K., Higaki, T., Isogai, Y., Nakano, T., Hasezawa, S., and Iba, K. (2016). Dominant and recessive mutations in the Raf-like kinase HT1 gene completely disrupt stomatal responses to CO2 in Arabidopsis. J. Exp. Bot. 67: 3251–3261.

Hayashi, M., Inoue, S., Ueno, Y., and Kinoshita, T. (2017). ARaf-like protein kinase BHP mediates blue light-dependent stomatal opening. Sci. Rep. 7: 45586.

Hõrak, H. et al. (2016). ADominant Mutation in the HT1 Kinase Uncovers Roles of MAP Kinases and GHR1 in CO2-Induced Stomatal Closure. Plant Cell 28: 2493–2509.

Hrabak, E.M. et al. (2003). The Arabidopsis CDPK-SnRK Superfamily of Protein Kinases. Plant Physiol. 132: 666–680.

Hwang, J.-U., Yim, S., Do, T.H.T., Kang, J., and Lee, Y. (2018). Arabidopsis thaliana Raf22 protein kinase maintains growth capacity during postgerminative growth arrest under stress. Plant Cell Environ. 41: 1565–1578.

Ichimura, K. et al. (2002). Mitogen-activated protein kinase cascades in plants: a new nomenclature. Trends Plant Sci. 7: 301–308.

Ishikawa, S., Barrero, J., Takahashi, F., Peck, S., Gubler, F., Shinozaki, K., and Umezawa, T. (2019a). Comparative phosphoproteomic analysis of barley embryos with different dormancy during imbibition. Int. J. Mol. Sci. 20: 451.

Ishikawa, S., Barrero, J.M., Takahashi, F., Nakagami, H., Peck, S.C., Gubler, F., Shinozaki, K., and Umezawa, T. (2019b). Comparative Phosphoproteomic Analysis Reveals a Decay of ABA Signaling in Barley Embryos during After-Ripening. Plant Cell Physiol.

Kodama, Y. (2016). Time Gating of Chloroplast Autofluorescence Allows Clearer Fluorescence Imaging In Planta. PLOS ONE 11: e0152484.

Lamberti, G., Gügel, I.L., Meurer, J., Soll, J., and Schwenkert, S. (2011). The Cytosolic Kinases STY8, STY17, and STY46 Are Involved in Chloroplast Differentiation in Arabidopsis. Plant Physiol. 157: 70–85.

Lee, S., Lee, M.H., Kim, J.-I., and Kim, S.Y. (2015). Arabidopsis Putative MAP Kinase Kinase Kinases Raf10 and Raf11 are Positive Regulators of Seed Dormancy and ABA Response. Plant Cell Physiol. 56: 84–97.

Matrosova, A., Bogireddi, H., Mateo-Peñas, A., Hashimoto-Sugimoto, M., Iba, K., Schroeder, J.I., and Israelsson-Nordström, M. (2015). The HT1 protein kinase is essential for red light-induced stomatal opening and genetically interacts with OST1 in red light and CO2-induced stomatal movement responses. New Phytol. 208: 1126–1137.

Nakagami, H., Sugiyama, N., Mochida, K., Daudi, A., Yoshida, Y., Toyoda, T., Tomita, M., Ishihama, Y., and Shirasu, K. (2010). Large-Scale Comparative Phosphoproteomics Identifies Conserved Phosphorylation Sites in Plants. Plant Physiol. 153: 1161–1174.

Nakashima, K., Fujita, Y., Kanamori, N., Katagiri, T., Umezawa, T., Kidokoro, S., Maruyama, K., Yoshida, T., Ishiyama, K., Kobayashi, M., Shinozaki, K., and Yamaguchi-Shinozaki, K. (2009). Three Arabidopsis SnRK2 Protein Kinases, SRK2D/SnRK2.2, SRK2E/SnRK2.6/OST1 and SRK2I/SnRK2.3, Involved in ABA Signaling are Essential for the Control of Seed Development and Dormancy. Plant Cell Physiol. 50: 1345–1363.

Nguyen, Q.T.C., Lee, S., Choi, S., Na, Y., Song, M., Hoang, Q.T.N., Sim, S.Y., Kim, M.-S., Kim, J.-I., Soh, M.-S., and Kim, S.Y. (2019). Arabidopsis Raf-Like Kinase Raf10 Is a Regulatory Component of Core ABA Signaling. Mol. Cells 42.

Nomoto, M. and Tada, Y. (2018). Cloning-free template DNApreparation for cell-free protein synthesis via two-step PCR using versatile primer designs with short 3′-UTR. Genes Cells 23: 46–53.

Ramachandiran, I., Vijayakumar, A., Ramya, V., and Rajasekharan, R. (2018). Arabidopsis serine/threonine/tyrosine protein kinase phosphorylates oil body proteins that regulate oil content in the seeds. Sci. Rep. 8: 1154.

Reddy, M.M. and Rajasekharan, R. (2006). Role of threonine residues in the regulation of manganese-dependent arabidopsis serine/threonine/tyrosine protein kinase activity. Arch. Biochem. Biophys. 455: 99–109.

Rudrabhatla, P., Reddy, M.M., and Rajasekharan, R. (2006). Genome-Wide Analysis and Experimentation of Plant Serine/ Threonine/Tyrosine-Specific Protein Kinases. Plant Mol. Biol. 60: 293–319.

Saruhashi, M., Ghosh, T.K., Arai, K., Ishizaki, Y., Hagiwara, K., Komatsu, K., Shiwa, Y., Izumikawa, K., Yoshikawa, H., Umezawa, T., Sakata, Y., and Takezawa, D. (2015). Plant Raf-like kinase integrates abscisic acid and hyperosmotic stress signaling upstreamof SNF1-related protein kinase2. Proc. Natl. Acad. Sci. 112: E6388–E6396.

Shinozaki, K., Yamaguchi-Shinozaki, K., and Seki, M. (2003). Regulatory network of gene expression in the drought and cold stress responses. Curr. Opin. Plant Biol. 6: 410–417.

Stevenson, S.R. et al. (2016). Genetic analysis of Physcomitrella patens identifies ABSCISIC ACID NON-RESPONSIVE (ANR), a regulator of ABA responses unique to basal land plants and required for desiccation tolerance. Plant Cell: tpc.00091.2016.

Sugiyama, N., Masuda, T., Shinoda, K., Nakamura, A., Tomita, M., and Ishihama, Y. (2007). Phosphopeptide Enrichment by Aliphatic Hydroxy Acid-modified Metal Oxide Chromatography for Nano-LC-MS/MS in Proteomics Applications. Mol. Cell. Proteomics 6: 1103–1109.

Tian, W. et al. (2015). A molecular pathway for CO2 response in Arabidopsis guard cells. Nat. Commun. 6: 6057.

Umezawa, T., Nakashima, K., Miyakawa, T., Kuromori, T., Tanokura, M., Shinozaki, K., and Yamaguchi-Shinozaki, K. (2010). Molecular Basis of the Core Regulatory Network in ABA Responses: Sensing, Signaling and Transport. Plant Cell Physiol. 51: 1821–1839.

Umezawa, T., Sugiyama, N., Mizoguchi, M., Hayashi, S., Myouga, F., Yamaguchi-Shinozaki, K., Ishihama, Y., Hirayama, T., and Shinozaki, K. (2009). Type 2C protein phosphatases directly regulate abscisic acid-activated protein kinases in Arabidopsis. Proc. Natl. Acad. Sci. 106: 17588–17593.

Umezawa, T., Sugiyama, N., Takahashi, F., Anderson, J.C., Ishihama, Y., Peck, S.C., and Shinozaki, K. (2013). Genetics and Phosphoproteomics Reveal a Protein Phosphorylation Network in the Abscisic Acid Signaling Pathway in Arabidopsis thaliana. Sci. Signal. 6: rs8–rs8.

Umezawa, T., Yoshida, R., Maruyama, K., Yamaguchi-Shinozaki, K., and Shinozaki, K. (2004). SRK2C, a SNF1-related protein kinase 2, improves drought tolerance by controlling stress-responsive gene expression in Arabidopsis thaliana. Proc. Natl. Acad. Sci. 101: 17306–17311.

Virk, N., Li, D., Tian, L., Huang, L., Hong, Y., Li, X., Zhang, Y., Liu, B., Zhang, H., and Song, F. (2015). Arabidopsis Raf-Like Mitogen-Activated Protein Kinase Kinase Kinase Gene Raf43 Is Required for Tolerance to Multiple Abiotic Stresses. PLOS ONE 10: e0133975.

Wagih, O., Sugiyama, N., Ishihama, Y., and Beltrao, P. (2016). Uncovering Phosphorylation-Based Specificities through Functional Interaction Networks. Mol. Cell. Proteomics 15: 236–245.

Wang, B., Liu, G., Zhang, J., Li, Y., Yang, H., and Ren, D. (2018). The RAF-like mitogen-activated protein kinase kinase kinases RAF22 and RAF28 are required for the regulation of embryogenesis in Arabidopsis. Plant J. 96: 734–747.

Wang, P., Xue, L., Batelli, G., Lee, S., Hou, Y.-J., Oosten, M.J.V., Zhang, H., Tao, W.A., and Zhu, J.-K. (2013). Quantitative phosphoproteomics identifies SnRK2 protein kinase substrates and reveals the effectors of abscisic acid action. Proc. Natl. Acad. Sci. 110: 11205–11210.

Yasumura, Y., Pierik, R., Kelly, S., Sakuta, M., Voesenek, L.A.C.J., and Harberd, N.P. (2015). An Ancestral Role for CONSTITUTIVE TRIPLE RESPONSE1 Proteins in Both Ethylene and Abscisic Acid Signaling. Plant Physiol. 169: 283–298.

Yoshida, R., Hobo, T., Ichimura, K., Mizoguchi, T., Takahashi, F., Aronso, J., Ecker, J.R., and Shinozaki, K. (2002). ABA-Activated SnRK2 Protein Kinase is Required for Dehydration Stress Signaling in Arabidopsis. Plant Cell Physiol. 43: 1473–1483.

Yoshida, T., Christmann, A., Yamaguchi-Shinozaki, K., Grill, E., and Fernie, A.R. (2019). Revisiting the Basal Role of ABA – Roles Outside of Stress. Trends Plant Sci. 24: 625–635.

